# Feral pig exclusion fencing provides limited fish conservation value on tropical floodplains

**DOI:** 10.1101/625053

**Authors:** N. J. Waltham, J. Schaffer

## Abstract

Efforts to protect and restore tropical wetlands impacted by feral pigs (*Sus scrofa*) in northern Australia have more recently included exclusion fences, an abatement response proposing fences improve wetland condition by protecting habitat for fish production and water quality. Here we tested: 1) whether the fish assemblage are similar in wetlands with and without fences; and 2) whether specific environmental processes influence fish composition differently between fenced and unfenced wetlands. Twenty-one floodplain and riverine wetlands in the Archer River catchment (Queensland) were surveyed during post-wet (June-August) and late-dry season (November-December) in 2016, 2017 and 2018, using a fyke soaked overnight (~14-15hrs). A total of 6,353 fish representing twenty-six species from 15 families were captured. There were no multivariate differences in fish assemblages between seasons, years and for fenced and unfenced wetlands (PERMANOVA, Pseduo-F <0.58, *P*<0.68). Late-dry season fish were considerably smaller compared to post-wet season: a strategy presumably to maximise rapid disposal following rain. At each wetland a calibrated Hydrolab was deployed (between 2-4 days, with 20min logging) in the epilimnion (0.2m), and revealed distinct diel water quality cycling of temperature, dissolved oxygen and pH (conductivity represented freshwater wetlands) which was more obvious in the late-dry season survey, because of extreme summer conditions. Water quality varied among wetlands, in terms of the daily amplitude, and extent of daily photosynthesis recovery, which highlights the need to consider local site conditions rather than applying general assumptions around water quality conditions for these types of wetlands examined here. Though many fish access (fenced and unfenced) wetlands during wet season connection, the seasonal effect of reduced water level conditions seems to be more over-improvised compared to whether fences are installed or not, as all wetlands supported few, juvenile, or no fish species because they had dried completed regardless of whether fences were present or not.

## Introduction

Wetlands (palustrine and lacustrine) that are located on floodplains away from riverine channels support rich aquatic plant and fauna communities [1–3]. However, some point after peak flood connection, aquatic organisms occupying these wetlands begin to face a moving land-water margin, until connection is broken, at which point the remaining wetland waterbodies typically support a non-random assortment of species, including fish [4, 5]. The duration, timing and frequency that off channel wetlands maintain lateral pulse connection to primary rivers is an important determining factor in broader contribution to coastal fisheries production [6–9]. In addition to connection, environmental conditions become important including water quality [10], access to shelter to escape predation and available food resources [11, 12]. Efforts by managers to restore wetland ecosystem values is increasing, nevertheless access to data establishing success of these programs are limited, which becomes important when attempting to establish biodiversity returns for the funding investment made by government or private investor organisations [13–15].

After floodplain wetlands begin receding and progressively disconnect from the main river channel, they become smaller and shallower [16] because of water loss via evaporation, groundwater recharge, or consumption by wildlife [17, 18]. In tropical north Australia, seasonal off channel wetlands are more pronounced owing to high evaporation rates, loss to groundwater [19], and in many situations waters quickly retract away from the banks and riparian shade [16]. At that point, it is thought that they become more prone to reduced water quality conditions - most notably reduced water depth [18], and high water temperatures [10, 20]. This increases aquatic fauna exposure risks to acute and chronic thresholds [21, 22]. In the late-dry season, fish confined to isolated wetlands on floodplains therefore have very limited avoidance options [10], unlike other fauna such as the freshwater crab also occupying seasonal tropical rivers in northern Australia, that will employ terrestrial re-location and access burrows when confronted with thermoregulation [23]. Fish must exploit available ephemeral aquatic habitats [24, 25], which can be specific to each wetland depending on orientation and location [26], depth and vegetation cover in the landscape [20], in order to survive until monsoonal rain reconnects overbank river networks again.

Across northern Australia, feral pigs (*Sus scrofa*) contribute wide scale negative impacts on wetland vegetation assemblages, water quality, biological communities and wider ecological processes [27, 28]. Feral pigs utilise an omnivorous diet supported by foraging or digging plant roots, bulbs and other below ground vegetation material over terrestrial or wetland areas [29]. This feeding strategy has a massive impact on wetland aquatic vegetation [30], which gives rise to soil erosion and benthic sediment re-suspension, reduced water clarity and eutrophication which becomes particularly critical late-dry season. The fact that limited data exits on the impact that feral pigs contribute to wetlands [31–34], places a strain on the ability for land managers to quantify the consequences of pig destruction [35]. Conversely, a lack of baseline data means quantifying success following expensive mitigation efforts is difficult.

Strategies focused on reducing or removing feral pigs from the landscape have been employed since their introduction to Australia [36], including poison baiting, aerial shooting, and trapping using specially constructed mesh cages [37]. Attempts to exclude feral pigs have also included installing exclusion fencing that border the wetland of interest. While advantages of installing fencing around wetlands has been examined only recently in Australia [32], those authors claim fencing might well be less effective particularly in situations where wetlands would normally dry during the dry season. Fencing is expensive to construct and maintain [37], but at the same time may prevent other non-target terrestrial fauna, such as kangaroos, from accessing wetlands which become particularly imperative late-dry season as regional water points [35]. Other terrestrial species including birds, snakes and lizards, for example, are generally able to access wetlands, though access for freshwater turtles might be hindered [38].

The aims were twofold: first to determine whether the model of non-randomness of fish stands here in wetlands, and secondly whether specific environmental conditions influence fish composition in wetlands with and without fences. These data are important and necessary given increasing government funding investment underway and planned in northern Australia for restoration of wetlands impacted by feral animals (including pigs) [10].

## Materials and methods

### Ethics Statement

This study was completed in accordance with the Queensland Animal Care and Protection Act 2001, and JCU animal ethics permit number A2178.

### Description of Study System

The Archer River catchment is located on Cape York Peninsula, north Queensland (Fig 1). The head waters of the river rise in the McIlwraith range on the eastern side Cape York, where it flows and then enters Archer Bay on the western side of the Gulf of Carpentaria; along with the Watson and Ward Rivers. The catchment area is approximately 13,820 km^2^, which includes approximately 4% (510 km^2^) of wetland habitats (https://wetlandinfo.des.qld.gov.au/wetlands/facts-maps/basin-archer/), such as estuarine mangroves, salt flats and saltmarshes, wet heath swamps, floodplain grass sedge, herb and tree Melaleuca spp. swamps and riverine habitat. The lower region of the catchment includes part of the Directory of Internationally Important Wetland network (i.e. nationally recognised status for conservation and cultural value) that extends along much of the eastern Gulf of Carpentaria, including the Archer Bay Aggregation, Northeast Karumba Plain Aggregation and Northern Holroyd Plain Aggregation. Two national parks are located in the catchment (KULLA (McIlwraith Range) National Park, and Oyala Thumotang National Park). Land use in the catchment is predominately grazing, with some mining activities planned in the next few years on the northern bank of the river (not within the area of this study).

**Fig 1.**
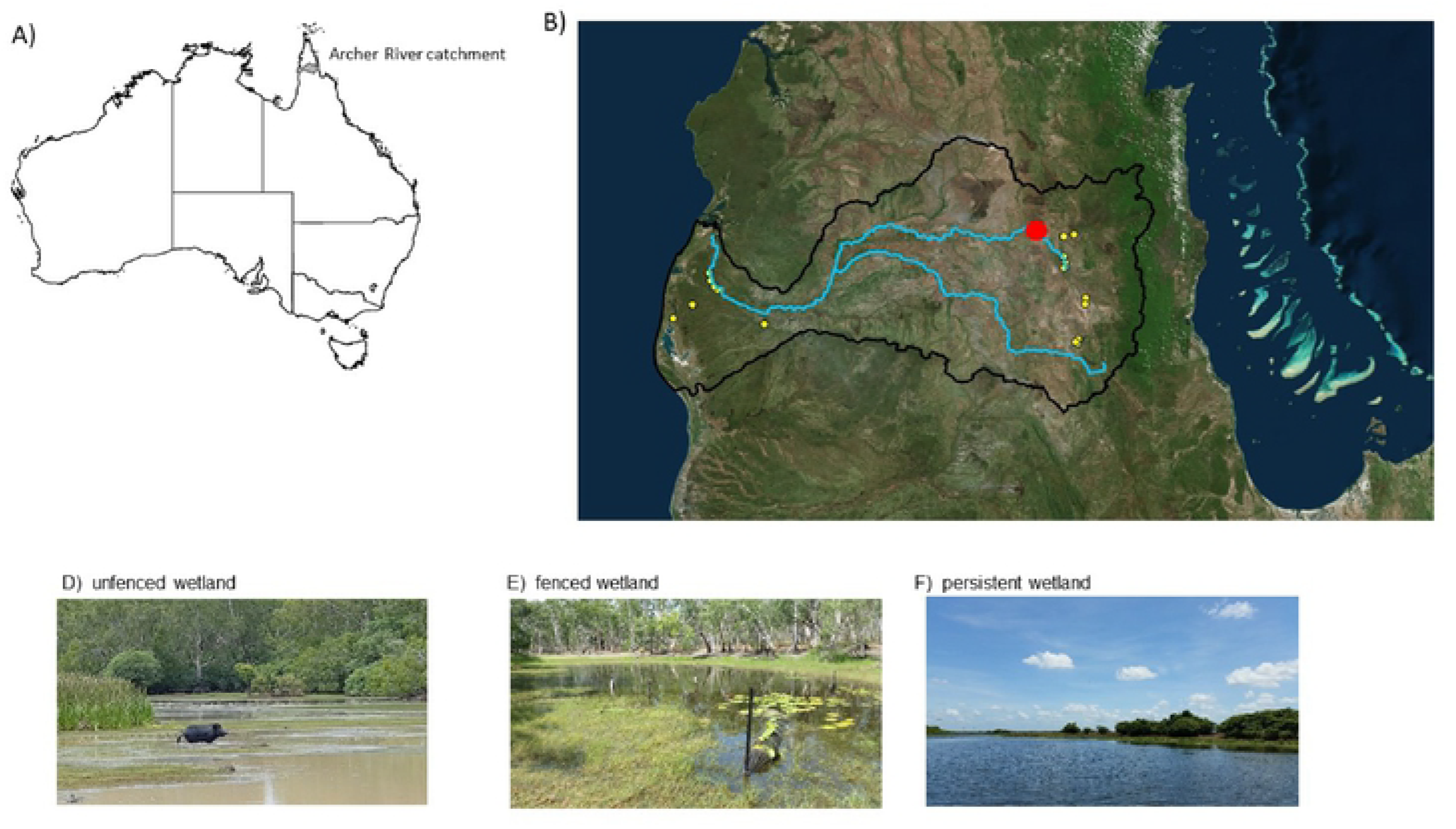
Location of wetlands in this study: a) location of the Archer River catchment in northern Queensland, Australia and b) wetland sites on the coastal floodplain and mid catchment where feral pig fencing has been completed around wetlands preventing access (yellow circles). The three wetland typologies (C - pig impacted wetlands that are shallow (typically <O.Sm deep), without submerged aquatic vegetation, turbid and eutrophic; D - fenced wetland preventing pig access that are deeper (typically <2m deep), clear with submerged aquatic vegetation present) exist across the catchment; and E - permanent wetlands that are deeper (typically <2m deep), steep sides limiting pig access, clear with submerged aquatic vegetation present Archer River gauge station (red circle).

Rainfall is tropical monsoonal with a strongly seasonal pattern where between 60% and 90% of total annual rain occurs between November and February. Long-term rainfall records for the catchment revealed highest wet season rainfall occurred in 1989/1999 (2515 mm), while the lowest was 1960/1961 (563.5 mm) [39]. Total antecedent rainfall for the wet season prior (Nov 2014 to Feb 2015) to this survey was 1081 mm, which is below the 10^th^ percentile for historical records. The wet seasons experienced through the years prior to this study (2010 to 2015) were among the wettest on record, within the 95^th^ percentile of the long-term data records. The low rainfall experienced during this study may have contributed to a short flood duration, and connection between study wetlands and the main Archer River, when compared to average or above average rainfall years (Fig 2).

**Fig 2.**
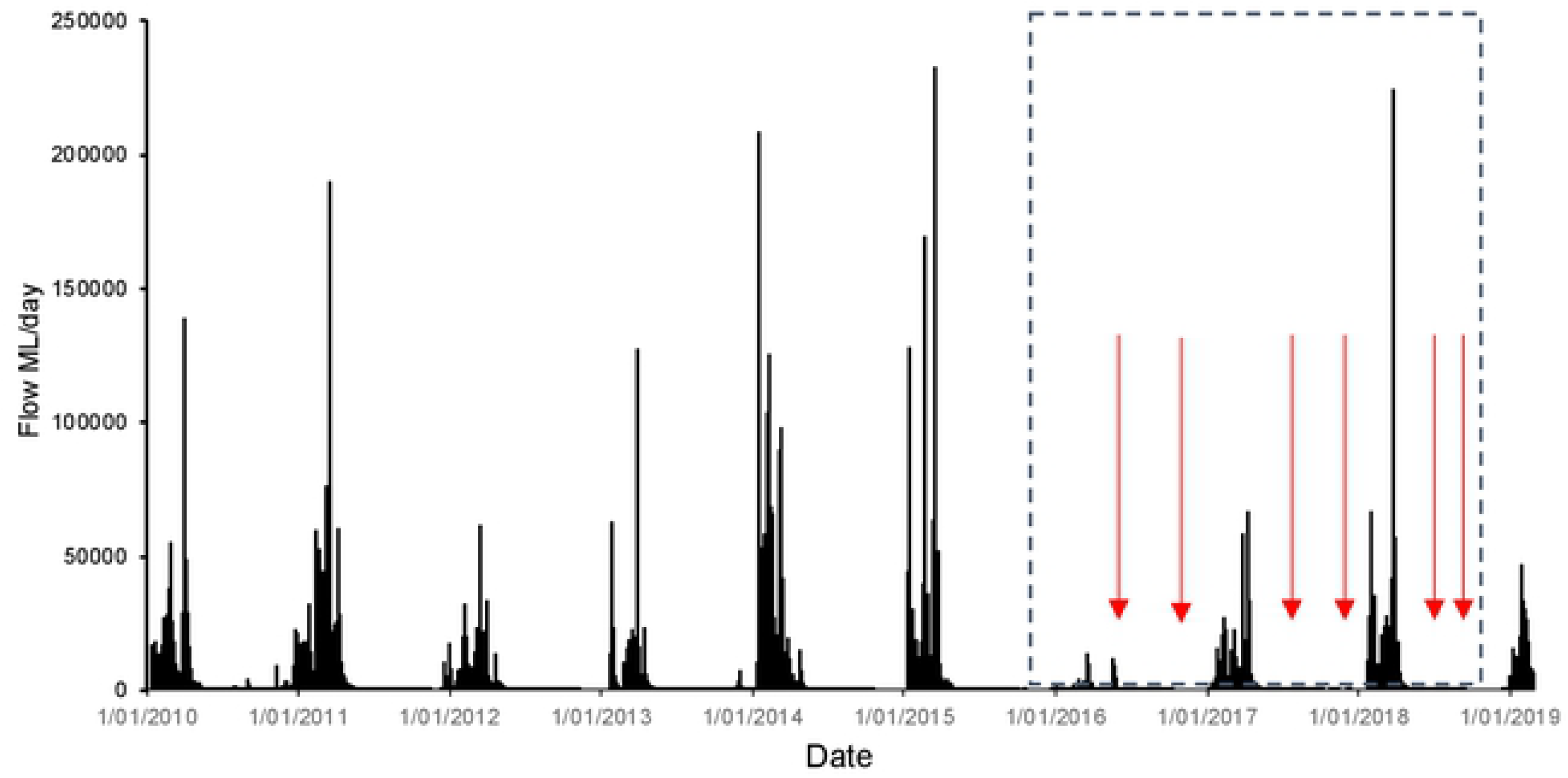
Daily discharge at the Archer River roadhouse gauge (Figure 1) before and during (dashed insert box) this study. Sampling occasions (arrows) are indicated. Data provided by the Queensland Government.

Twenty-one wetlands from the Archer River catchment were sampled for this project. These included both floodplain and riverine wetlands that were not on the main flow channels, but rather were on anabranches and flood channels that connect to the main channels only during high flow conditions. All wetlands have been historically damaged by pigs (and cattle to a lesser extent) for up to 160 years [40, 41], until recently, where a small number were fenced to abate feral pig and/or cattle from accessing wetlands, in accordance with the feral animal research and management agenda (to meet the objectives of traditional owners in the region) of both Kalan enterprises and Aak Puul Ngangtam, and their partner.

The characteristics of each wetland are summarised in Table S1. Here, sampling focused on two periods: 1) immediately following the wet season after disconnection between the river and wetlands (hereafter referred to as post wet season); and 2) late-dry season (hereafter late-dry) in 2016, 2017 and 2018. Each sampling campaign was completed over 14 days with six campaigns in total (post-wet and late-dry season in 2016, 2017 and 2018).

### Field Methods

In each wetland, a calibrated high frequency Hydrolab multi-parameter logger (OTT Hydromet USA) was deployed (0.2m depth) for between 2 and 4 days to record epilimnion (0.2m) water temperature, dissolved oxygen (%), electrical conductivity and pH every 20mins; logging at this frequency provides explicit insight into diel changes in environmental water processes [20, 22]. Weather conditions were fine with wetlands surveyed on the falling limb of the hydrograph.

Fish were collected in wetlands using a fyke net (0.8m opening, double 4m wing panels, 1mm stretch mesh) that was soaked overnight (approximately 14:00 to 09:00). Wetlands substantially impacted by feral pigs; secchi disk depth < 0.1m, no submerged or floating aquatic plants exist, while the fenced wetlands were generally deeper (up to 1.5m), and had submerged aquatic vegetation (Fig. 1). Fish were placed in a tub (~150L) temporarily, identified, measured (standard length, mm) and returned to the wetland alive in accordance with Australian laws (except for a small number that were kept food web studies, not shown here, under Australian law).

### Data Analysis

There are two main biases in the sampling method here: 1) that the sampling technique potentially capture large numbers of schooling fish along the wetland margins; and 2) the fact that predatory aquatic fauna including fish, snakes(macleays watersnakes (*Pseudoferania polylepis*), file snakes (*Acrochordus arafurae*)) and freshwater turtles (*Chelodina oblonga*, *Chelodina canni* and *Emydura s. worrelli*), were periodically trapped in the net for hours means that they could consume fish trapped in nets. To overcome these uncertainties, analyses were based on presence/absence of species. Presence/absence provide robust data when relative abundance are of doubtful validity because they treat species with a diversity of behaviours, trophic functions, and spatial distribute in a more equivalent way than fully quantitative techniques [42].

Multivariate differences were examined using PERMANOVA (Anderson, 2001), using the Bray-Curtis similarities measure [43] with significance determined from 10,000 permutations of presence/absence transformation. Multivariate dispersion were tested using PERMDISP, however, homogeneity of variance could not be stabilized with transformation, and therefore untransformed data were used. Three factors where included: years (fixed), season (fixed); and fenced/unfenced (random). These factors were determined *a-prior* during study design.

Spatial patterns in multivariate fish assemblage structure and the importance of explanatory data sets were analysed using a multivariate classification and regression tree (mCARTs) [44] package in R (version 3.4.4). Analysis was conducted using presence/absence transformed fish data for the 10 species that occurred in >20% of wetland sites (to remove rare species from this analysis). Selection of the final tree model was conducted using 10-fold cross validation, with a 1-SE tree; the smallest tree with cross validation error within 1 SE of the tree with the minimum cross validation error [45]. The relative importance of the explanatory variables were assessed to determine those with a high overall contribution to tree node split, with the best overall classifier being given a relative importance of 100%.

Kolmogorov-Smirnov (K-S) two-sample tests determined differences in the overall shape of fish body size distribution using a Bonferroni correction for multiple comparisons. K-S tests take into account differences between the location, skew, and kurtosis of frequency distributions; but do not identify which of these parameters are driving distributional differences. Therefore, we report the following characteristics of each body size distribution to further describe any differences found: mean, standard deviation (sd), minimum value (min), maximum value (max), the range of values, skewness, and kurtosis.

## Results

### Hydrology and Wetland Water Quality

Wet season rainfall totals in the Archer River catchment were low during the study period compared to the preceding years (Fig. 2), with rainfall within the 10^th^ percentile for historical recordings held by the Australian Bureau of Meteorology. This means that some caution is necessary with interpretation of these data; namely that floodplain connectivity under higher rainfall years is likely to have a longer duration when compared to lower connection duration under the current rainfall conditions.

A full summary of water quality data are provided in Supplementary files (S1). In summary, water temperatures during the study period were generally about 26°C (Table 1). Minimum water temperature recordings as low as 18°C, while maximum temperatures occurred in November 2016 survey reached above 40°C for several hours of the day in some instances. The water column exhibited pronounced diel temperature periodicity; one or two hours after sunrise each day. Near-surface water temperatures began to rise at an almost linear rate for a period of 8.0 ± 0.5 hours, generally reaching daily maxima during the middle of the afternoon. The mean daily temperature amplitude was 6.2ºC (highest daily amplitude 9.6ºC, lowest 4.4°C). For the remaining 16 hours of each day, near-surface water temperatures gradually declined reaching minimum conditions shortly after sunset.

The electrical conductivity (EC) was very low (Table S1) during the post wet season surveys, while the late-dry season conductivity was generally higher in wetlands, a consequence of evapo-concentration. The lowest wetland in the catchment (AR08 located on the coastal floodplain) recorded the highest conductivity, suggesting connection with tidal water from the nearby estuary at some stage.

There was evidence of cyclical daily DO fluctuations supporting the contention that biological diel periodicity processes were probably not significantly inhibited in all wetlands (Fig. 3). Daily minimum DO concentrations were low enough to suggest there was enough respiratory oxygen consumption to measurably affect water quality, particularly so at the pig impacted wetlands, but also during the late-dry season survey in November 2016. Dissolved oxygen (DO) seemed to reach daily minima conditions, well below the asphyxiation thresholds of sensitive fish species, in the early morning hours during all surveys. In the examples shown, after the morning low DO, conditions generally recovered to approximately 50%, but reaching a high of 100-160% in the late afternoon (before sunset).

**Fig 3.**
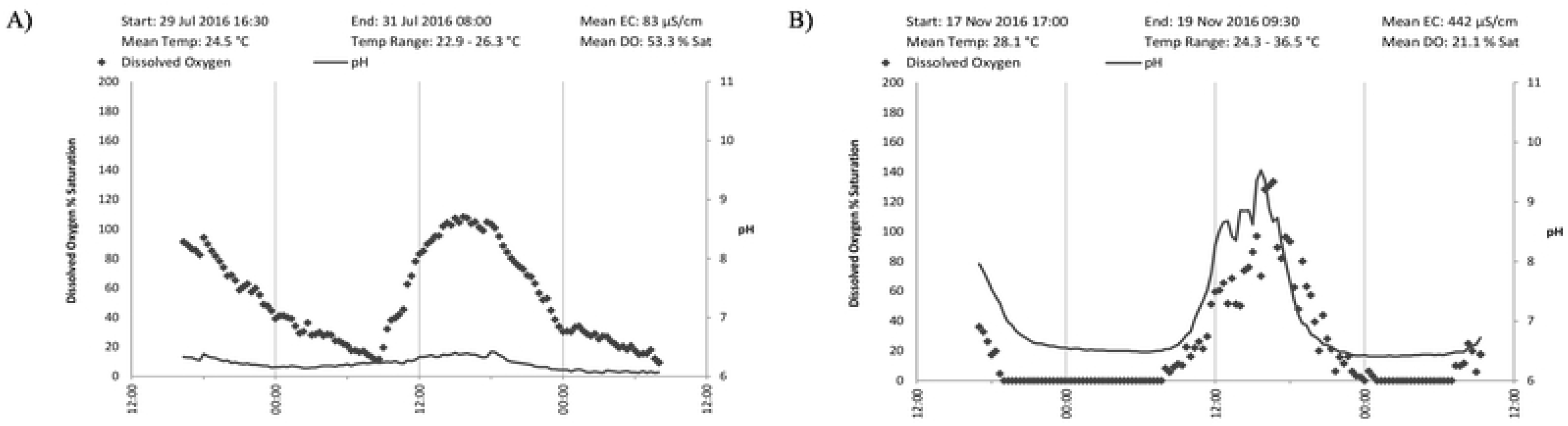
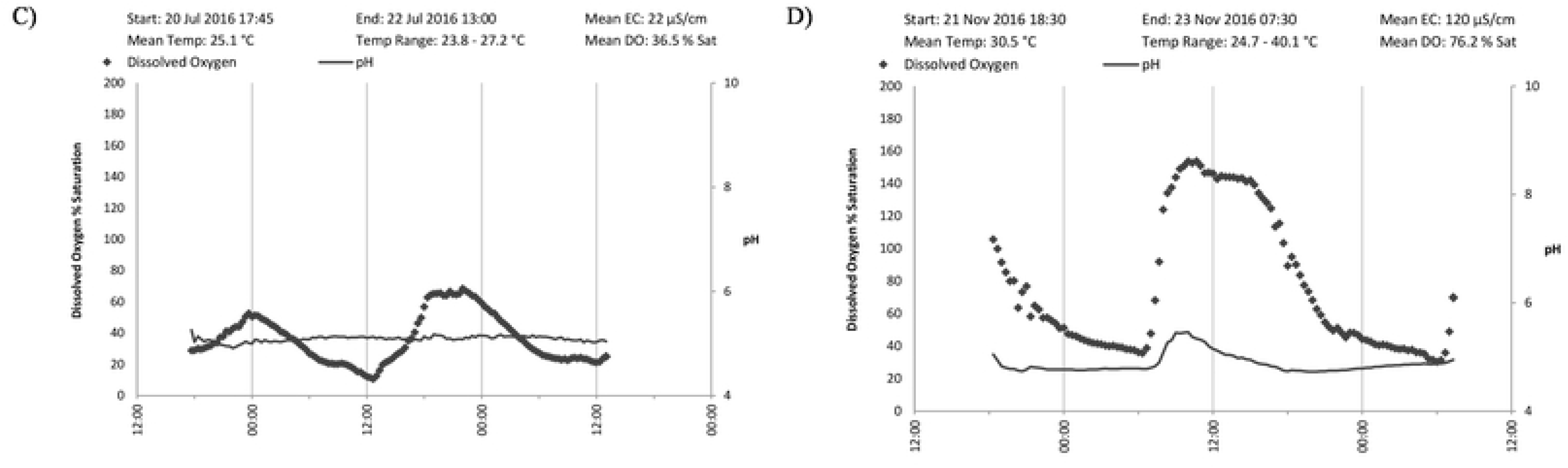
Examples of the diel dissolved oxygen, pH, water temperature and conductivity cycling in Archer River wetlands. These examples are from KA06 during post-wet season (a), and late-dry season (b) in 2016, and AROJ during post wet season (c) and late-dry season (d) in 2016.

**Fig 4.**
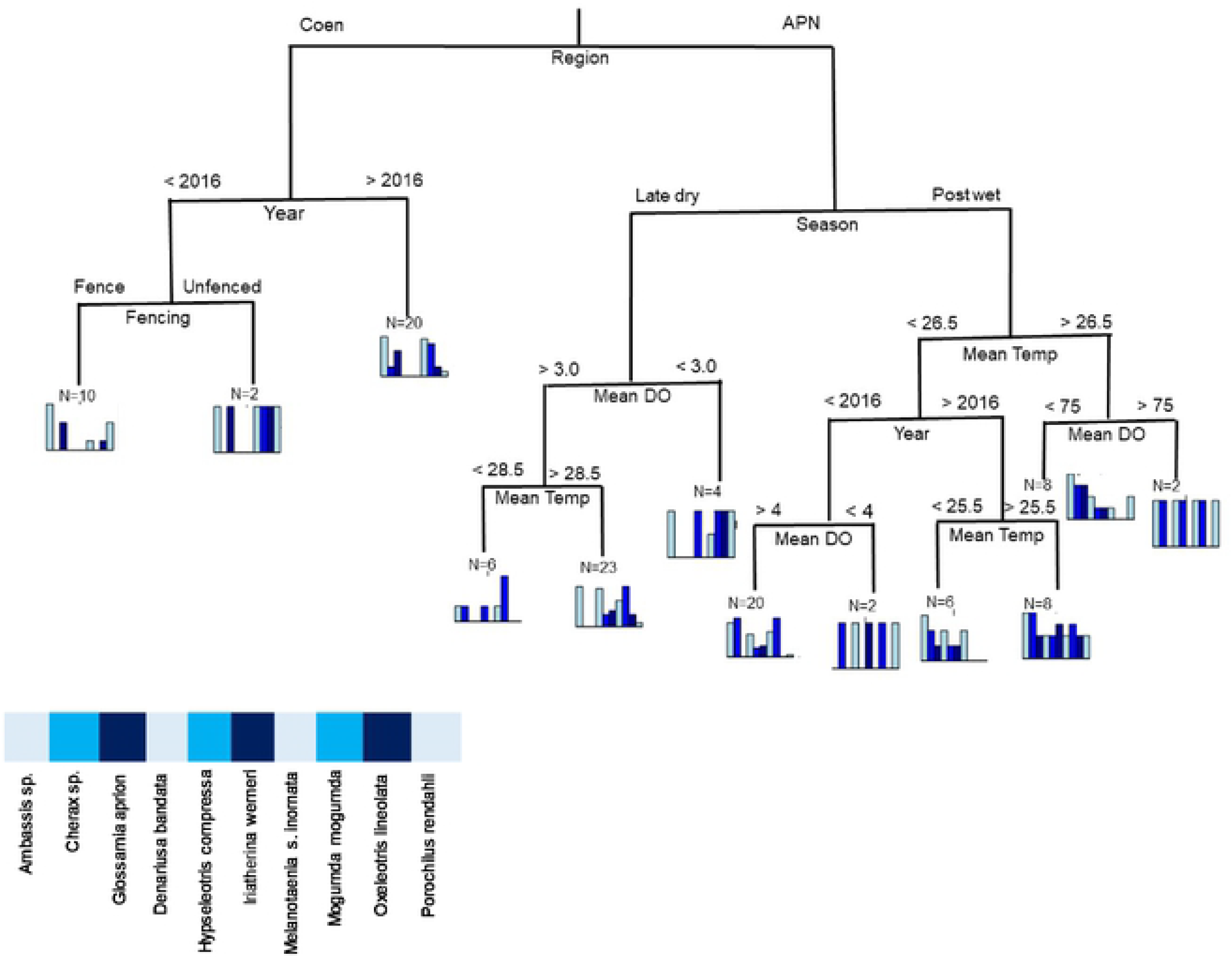
Multivariate regression tree showing the major divisions in the database on assemblage composition. Each of the splits are labelled with the contributing variable, and the division threshold (in the case of electronic conductivity; EC, and dissolved oxygen; DO). The length of the descending branches is proportional to the divergence between groups. Bar plots represent the fish assemblage composition at the corresponding colour code node sharing the same attributes. Values in the bar plots represent the relative frequencies of occurrence of each taxon within a same node.

pH is also potentially subject to the same kinds of biogenic fluctuations as DO, due to consumption of carbon dioxide (i.e., carbonic acid) by aquatic plants and algae during the day (through photosynthesis), and net production of carbon dioxide at night. If respiratory oxygen consumption is predominant, DO concentrations are low and pH values are generally moderately acidic to neutral, which was the case for wetlands examined here. All photosynthetically active organisms utilise carbon dioxide as a preferred carbon source. Some species (including most green algae) are unable to photosynthesise if carbon dioxide is unavailable, but there are other species (including most cyanobacteria and submerged macrophytes) which can utilise bicarbonate as an alternative carbon source. Carbon dioxide consumption causes pH to rise to values in the order of 8.6 to 8.7 (but that was not the case here during this survey period).

### Fish Community

A total of 6,353 fish were captured, representing twenty-six species from 15 families (Table 1). The most common species was the freshwater glassfish (*Ambassis sp.*, 51% total catch), delicate blue-eyes (*Pseudomugil tenellus*, 11%), and northern purple-spot gudgeon (*Morgunda morgunda*, 9%). A greater number of fish species were caught in the post wet season survey, with a lower number captured during the late-dry season, including the northern purple-spot gudgeon (*Morgunda mogunda*), chequered rainbow fish (*Melanotaenia s. inornata*), and the empire gudgeon (*Hypseloptris compressa*). In addition to fish, we captured a freshwater crayfish (*Cherax sp*.), macleays watersnakes (*Pseudoferania polylepis*) and freshwater turtles (*Chelodina oblonga* and *Emydura s. worrelli*) in most wetlands, notably during post wet season. Overall, there was no significant difference between seasons, fenced/unfenced wetlands and among years (PERMANOVA, Pseduo-F <0.58, *P*<0.68).

With a reduced list confined to dominant species, occurrence profiles for groups in the terminal branches of the mCART analysis (Fig. 5) show two initial wetland groups based on a split supported by region, with wetlands in the Coen (upper catchment) region separating from those wetlands in the coastal plains. Following the left branch there is inter-annual variation among wetlands, and a second terminal node based on whether wetlands were fenced in 2016, but not so in 2017 and 2018 data. Following the right branch (APN, coastal plains), the first node separates seasons, and following late-dry season wetlands further separate based on mean dissolved oxygen (~3.0%), and then mean temperature (~28.5°C). The post-wet season branch appears to have more separation among data, with a separation based on mean water temperature (~26.5°C), years, and then finally dissolved oxygen (~4%).

**Fig 5.**
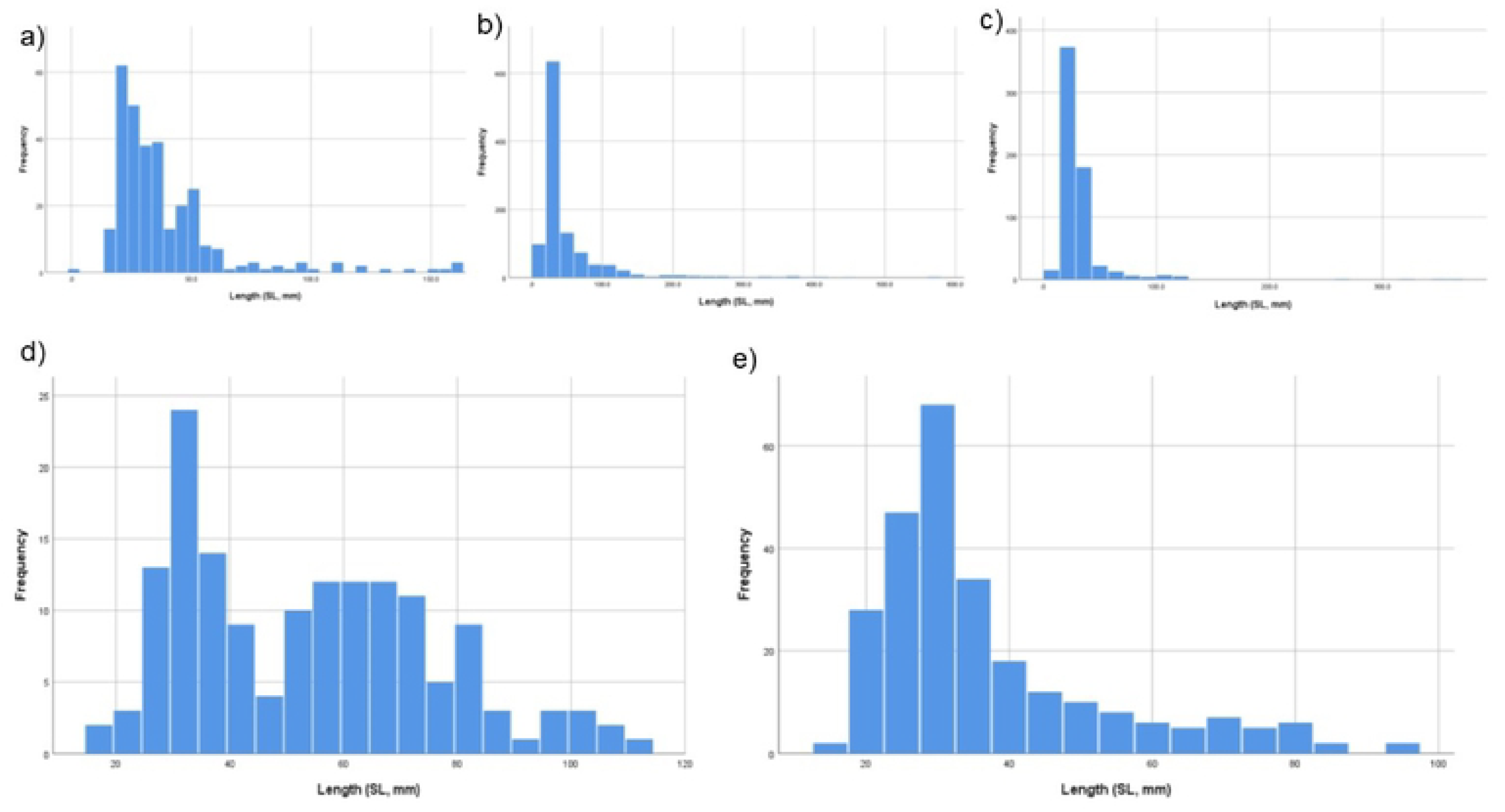
Pooled fish length (standard length, mm) frequency comparison of(a) 2016, (b) 2017 and (c) 2018. Comparison for *Mogurnda mogurnda* (d) post wet season; and (e) late-dry season (years combined).

Mean fish body size distributions differed between the three sample years (with fish for each wetland and survey pooled) (KS, *P* < 0.001, Table S2 – S5), with larger fish measured in 2017 (50.5mm) compared to 2016 (38.7mm) and 2018 (31.6mm), despite the assemblages having similar size ranges. When comparing the overall fish size distribution by pooling years, post wet season fish were larger (44.9mm) when compared to fish in the late-dry season (39.7mm) (KS, *P* < 0.01). For some fish species such as the chequered rainbow fish (*Melanotaenia s. inornata*), the post wet season (32.5mm) was similar when compared to late-dry season (38.4mm) (KS, *P* = 0.06, S3). In contrast, the northern purple-spot gudgeon (*Mogurnda mogurnda*) was larger post-wet season (52.8mm) compared to late-dry season (37.1mm) (KS, *P* < 0.01, Figure 5, S4).

## Discussion

While installation of fences can protect terrestrial ecosystem services from feral impacts [46], however, in the case here abatement fences appear to offer little over-improvised fish additional value compared to those that are not fenced. Many fish indeed access both fenced and unfenced wetlands during wet season connection, however, the seasonal effects of reduced water level conditions and the loss of fish assemblage as the dry season progresses is a pattern that remained regardless of fencing. To this end, installation of expensive exclusion fences might not offer additional protection to fish species occupying these tropical floodplain wetlands. The same conclusion was reported by [32] where those authors surveyed strongly seasonal wetlands (similar to the wetlands here) elsewhere in northern Australia, and concluded that the seasonal dry down of wetlands ultimately prohibits the wetland contribution to future year successful fish recruitment.

The low species richness in wetlands relative to the main Archer River channel might be a consequence of the frequency and duration of connection between wetlands and the main Archer river channel. The wet season rainfall immediately prior, and during this survey, was within the 10^th^ percentile for historical records. In research elsewhere, a longer connection duration was shown to result in more fish present post wet season, and conceivably more species present late-dry season [6, 47]. Examples exist where longer connection between main river channels and wetlands contributes positively to fish growth rates and higher abundance and diversity of fish [24, 26, 48]. It is also possible that the field methods used here confound our ability to determine the full species composition in wetlands – this could be overcome by using additional survey techniques, including multi-panel gill nets, traps or electrofishing (though we attempted to electrofish these wetlands however, conductivity was too low to effectively use that method).

An obvious characteristic of the fish data were larger, presumably adult, individuals following the disconnection of wetlands after the wet season compared to small individuals present in the late-dry season. On this basis, it is possible that the wetlands serve as important refugia for successful recruitment of freshwater fish, that adult fish remaining in the wetlands after disconnection are able to complete important life cycle stages. The fact that we did not catch large fish in the late-dry season suggests that adult fish might be lost as the dry season progress, consumed either by larger predators such as estuarine crocodiles (*Crocodylus porosus*), or birds feeding in the shallow waters. Wetlands are also popular feeding and roosting locations for birds [2, 49]; we observed a large number of birds at most wetlands in the late-dry season. The value of wetlands to wader birds is limited by the condition of wetlands [50, 51], but wetlands provide an important nutrient subsidy more broadly on seasonal floodplains [52, 53]. Hurd et al. (2017) postulates that differences in fish communities among off channel waters could be more influenced by the presence of piscivorous predators which reduce or eliminate prey species, or even via a function of competitive exclusion may occur within fish guilds as resources diminish in the late-dry season. Examining this point could be achieved by investigating the species niche width [54, 55] in drying waters by constructing food webs in individual waters to determine species ranges and changes with fencing treatment and comparing post wet season and late-dry season conditions.

In the late-dry season for the few fish species present, juveniles dominated the catch regardless whether wetlands were fenced. Having small recruits in the late dry period might be an important strategy in maximising, quick, dispersal after connectivity with the onset of the wet season [56]. Moreover, late season conditions with no flow and warm conditions might favour larval development [57, 58]. *Melanotaeniid* rainbowfish, for example, have a flexible reproductive behaviour that is well adapt to deal with the vagaries of temporal variation in habitat conditions [59]. The same is true for both *Eleotrid gudgeon* species found here, with smaller recruits presumably ready for wide-scape distribution with the onset of seasonal flow. Pusey et al. (2018) provides a case that the reproduction success of freshwater fish in northern Australia could in fact hinge on antecedent flow patterns across the landscape, and that this flexibility ensures population level success [60]. This production strategy might be particularly pertinent given the below average summer rainfall totals witnessed during this survey, particularly when compared to previous years.

As the dry season takes hold, water quality conditions progressively deteriorate owing mostly to increasing impact from rooting pigs accessing wetland vegetation. Generally, fenced wetlands change little in terms of water conditions (Figure 6). However, it is the late-dry season when water conditions are poorest and therefore most critical to fish. Unfenced wetlands tended to be shallower, highly turbid, and most notably experience water temperatures that exceed acute thermal thresholds for fish [31]. The solubility of dissolved oxygen in water is strongly affected by temperature (i.e. high temperature reduces dissolved oxygen solubility [61]. Data on hypoxia tolerances of local freshwater fish species in northern Queensland is available [62], and while tolerances vary between species and life stages, there were obvious periods in wetlands when these threshold limits are exceeded. During the critical periods, fish must regulate breathing either via increasing ventilation rates [63], or by rising to the surface to utilise aquatic surface respiration and/or air gulping (e.g. tarpon, *Megalops cyprinoides*). In any case, the capacity for fish to do that safely depends on the timing of the oxygen sag and antecedent conditions, though notably it appears that most of the hypoxia-induced fish kills is actually due to exposure (e.g., thermal stress and sunburn) resulting from the animals’ need to remain at the surface during the heat of the day in order to access available oxygen for respiration. Increasing these risks to fish can have important chronic effects including reducing physical fitness of fish to successfully contribute to future populations [64, 65].

**Fig 6.**
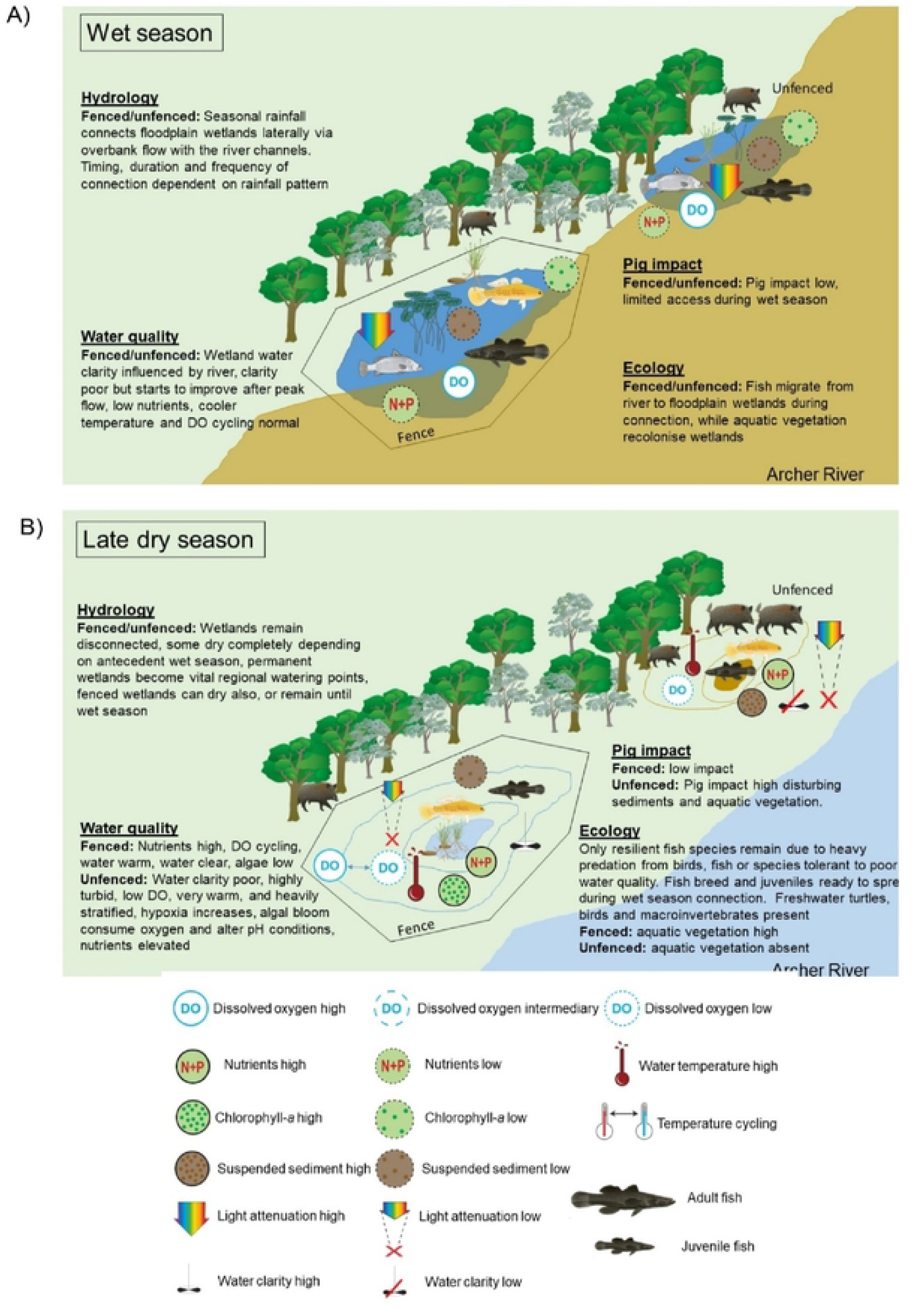
Conceptual diagram of wetland ecosystem conditions during (a) wet season, and (b) late-dry season. During the wet season, the latera.1 connection between the Archer River channel and wetlands occurs, during which fish can access wetlands and water quality is generally best because feral pig impact is minimal regardless of fencing. The dry season results in water retracting from the land margins, allowing pigs to access unfenced wetlands. At this stage, water quality conditions are poor in unfenced wetlands with high turbidity/nutrients and temperature, and dissolved oxygen is generally critical for fish. Fenced wetlands become shallower too, through temperature and dissolved oxygen cycling reduced, turbidity is low, while nutrients can be also high. Regardless of fencing, fish community reduced to a few resilient species dominated by juveniles ready for rapid dispersal when wet seasons commences again.

The cultural and ecological value of coastal wetlands means that management intervention is increasingly necessary to ensure they remain productive and viable habitat (Creighton *et al.*, 2015). These data support a model that damage to wetlands from pig activities not only contributes to reduced aquatic habitat, through loss of aquatic vegetation communities, but also probably has secondary impacts including water temperature and asphyxiation risks for many hours each day, that are higher than when compared to fenced wetlands (Figure 6). However, fish occupying fenced and unfenced wetlands here were similar, particularly in the late-dry season where those remaining few species were juveniles ready for wet season re-distribution. On this basis, installing fences to both floodplain and riverine wetlands that were not on the main flow channels, but rather were on anabranches and flood channels that connect to the main channels only during high flow conditions, seems to offer little additional habitat value for fish. Where wetlands are largely ephemeral and will dry anyway, or where wetlands remain until the next seasons rain connection; species abundance and/or diversity is not improved by restricting feral pig access. Further research is necessary to examine climate change resilience on permanent wetlands (and managed wetlands) particularly whether they provide a similar level of refugia [66].

## Funding

This study is supported by the National Environment Science Program (Northern Australian Hub) awarded to CSIRO, James Cook University, and the Queensland Government.

## Acknowledgements

This project builds on a long-term feral animal management and monitoring program developed by Kalan enterprises and Aak Puul Ngangtam (APN) and their partners. Kalan and APN have developed their feral animal research and management agenda to meet the objectives of traditional owners in the region and have invited science organisations (CSIRO, James Cook University and the Department of Science and Environment) to contribute to the outcomes. APN and Kalan have conducted systematic feral pig control and monitoring in the Archer River basin for the past 6 years. We thank the reviewers who improved this manuscript considerably.

## Author Contributions

NW conceived and designed the study, NW performed the data analysis. NW and JS completed fieldwork and prepared the manuscript.

## References

1. Ambrose RF, Meffert DJ. Fish-assemblage dynamics in Malibu lagoon, a small, hydrologically altered estuaryin southern California. Wetlands. 1999;19(2):327–40.

2. Brandolin PG, Blendinger PG. Effect of habitat and landscape structure on waterbird abundance in wetlands of central Argentina. Wetlands ecology and management. 2016;24(1):93–105.

3. Jiang T-t, Pan J-f, Pu X-M, Wang B, Pan J-J. Current status of coastal wetlands in China: degradation, restoration, and future management. Estuarine, Coastal and Shelf Science. 2015;164:265–75.

4. Arrington DA, Winemiller KO. Habitat affinity, the seasonal flood pulse, and community assembly in the littoral zone of a Neotropical floodplain river. Journal of the North American Benthological Society. 2006;25(1):126–41.

5. Pander J, Mueller M, Geist J. Habitat diversity and connectivity govern the conservation value of restored aquatic floodplain habitats. Biological Conservation. 2018;217:1–10.

6. Hurd LE, Sousa RG, Siqueira-Souza FK, Cooper GJ, Kahn JR, Freitas CE. Amazon floodplain fish communities: Habitat connectivity and conservation in a rapidly deteriorating environment. Biological Conservation. 2016;195:118–27.

7. Bennett MG, Kozak JP. Spatial and temporal patterns in fish community structure and abundance in the largest US river swamp, the Atchafalaya River floodplain, Louisiana. Ecology of freshwater fish. 2016;25(4):577–89.

8. Galib SM, Lucas MC, Chaki N, Fahad FH, Mohsin A. Is current floodplain management a cause for concern for fish and bird conservation in Bangladesh’s largest wetland? Aquatic Conservation: Marine and Freshwater Ecosystems. 2018.

9. Górski K, De Leeuw J, Winter H, Khoruzhaya V, Boldyrev V, Vekhov D, et al. The importance of flooded terrestrial habitats for larval fish in a semi-natural large floodplain (Volga, Russian Federation). Inland Waters. 2016;6(1):105–10.

10. Waltham N, Schaffer J. Thermal and asphyxia exposure risk to freshwater fish in feral-pig-damaged tropical wetlands. Journal of fish biology. 2018.

11. Blanchette ML, Davis AM, Jardine TD, Pearson RG. Omnivory and opportunism characterize food webs in a large dry-tropics river system. Freshwater Science. 2014;33(1):142–58.

12. Jardine TD, Pettit NE, Warfe DM, Pusey BJ, Ward DP, Douglas MM, et al. Consumer-resource coupling in wet-dry tropical rivers. J Anim Ecol. 2012;81(2):310–22. Epub 2011/11/23. doi: 10.1111/j.1365-2656.2011.01925.x. PubMed PMID: 22103689.

13. Waltham N, Fixler S. Aerial Herbicide Spray to Control Invasive Water Hyacinth (Eichhornia crassipes): Water Quality Concerns Fronting Fish Occupying a Tropical Floodplain Wetland. Tropical Conservation Science. 2017;10:1940082917741592.

14. Weinstein MP, Litvin SY. Macro-Restoration of Tidal Wetlands: A Whole Estuary Approach. Ecological Restoration. 2016;34(1):27–38.

15. Zedler JB. What’s New in Adaptive Management and Restoration of Coasts and Estuaries? Estuaries and Coasts. 2016:1–21.

16. Pusey BJ, Arthington AH. Importance of the riparian zone to the conservation and management of freshwater fish: a review. Marine and Freshwater Research. 2003;54:1–16.

17. McJannet D, Marvanek S, Kinsey-Henderson A, Petheram C, Wallace J. Persistence of in-stream waterholes in ephemeral rivers of tropical northern Australia and potential impacts of climate change. Marine and Freshwater Research. 2014.

18. Pettit N, Jardine T, Hamilton S, Sinnamon V, Valdez D, Davies P, et al. Seasonal changes in water quality and macrophytes and the impact of cattle on tropical floodplain waterholes. Marine and Freshwater Research. 2012;63(9):788–800.

19. Petheram C, McMahon TA, Peel MC. Flow characteristics of rivers in northern Australia: implications for development. Journal of Hydrology. 2008;357(1):93–111.

20. Wallace J, Waltham N, Burrows D. A comparison of temperature regimes in dry-season waterholes in the Flinders and Gilbert catchments in northern Australia. Marine and Freshwater Research. 2017;68(4):650–67.

21. Burrows D, Butler B. Priminary studies of temperature regimes and temperature tolerance of aquatic fauna in freshwater habitats of northern Australia. Australian Centre of Tropical Freshwater Research (12/01): James Cook University, Townsville, 2012 Contract No.: 12/01.

22. Wallace J, Waltham NJ, Burrows DW, McJannet D. The temperature regimes of dry-season waterholes in tropical northern Australia: potential effects on fish refugia. Freshwater Science. 2015;34(2):663–78.

23. Waltham NJ. Acute thermal effects in an inland freshwater crab Austrothelphusa transversa (von Martens, 1868) occupying seasonal, tropical rivers. Journal of Crustacean Biology. 2018.

24. Love S, Phelps Q, Tripp S, Herzog D. The importance of shallow-low velocity habitats to juvenile fish in the middle Mississippi River. River Research and Applications. 2017;33(3):321–7.

25. Phelps QE, Tripp SJ, Herzog DP, Garvey JE. Temporary connectivity: the relative benefits of large river floodplain inundation in the lower Mississippi River. Restoration Ecology. 2015;23(1):53–6.

26. Schomaker C, Wolter C. The contribution of long-term isolated water bodies to floodplain fish diversity. Freshwater Biology. 2011;56(8):1469–80.

27. Baber DW, Coblentz BE. Density, home range, habitat use, and reproduction in feral pigs on Santa Catalina Island. Journal of Mammalogy. 1986;67(3):512–25.

28. Krull CR, Choquenot D, Burns BR, Stanley MC. Feral pigs in a temperate rainforest ecosystem: disturbance and ecological impacts. Biological invasions. 2013;15(10):2193–204.

29. Ballari SA, Barrios-García MN. A review of wild boar S us scrofa diet and factors affecting food selection in native and introduced ranges. Mammal Review. 2014;44(2):124–34.

30. Doupé RG, Mitchell J, Knott MJ, Davis AM, Lymbery AJ. Efficacy of exclusion fencing to protect ephemeral floodplain lagoon habitats from feral pigs (Sus scrofa). Wetlands Ecology and Management. 2010;18(1):69–78.

31. Waltham NJ, Schaffer JR. Thermal and asphyxia exposure risk to freshwater fish in feral-pig-damaged tropical wetlands. Journal of Fish Biology. 2018;93(4):723–8.

32. Doupe RG, Mitchell J, Knott MJ, Davis AM, Lymbery AJ. Efficacy of exclusion fencing to protect ephemeral floodplain lagoon habitats from feral pigs (Sus scrofa). Wetlands Ecology Management. 2010;18:69–78.

33. Mitchell J, Mayer R. Diggings by feral pigs within the Wet Tropics World Heritage Area of north Queensland. Wildlife Research. 1997;24(5):591–601.

34. Steward AL, Negus P, Marshall JC, Clifford SE, Dent C. Assessing the ecological health of rivers when they are dry. Ecological Indicators. 2018;85:537–47.

35. Australia Co. Threat abatement plan for predation, habitat degradation, competition and disease transmission by feral pigs (Sus scrofa) (2017) — Background Document. Canberra: Department of Environment and Energy, 2017.

36. Fordham D, Georges A, Corey B, Brook BW. Feral pig predation threatens the indigenous harvest and local persistence of snake-necked turtles in northern Australia. Biological Conservation. 2006;133(3):379–88.

37. Ross B, Waltham NJ, Schaffer J, Jaffer T, Whyte S, Perry J, et al. Improving biodiversity outcomes and carbon reduction through feral pig abatement. Cairns: Balkanu Cape York Development Corporation Ltd Pty, 2017.

38. Waltham N, Schaffer J. Continuing aquatic assessment of wetlands with and without feral pig and cattle fence exclusion, Archer River catchment. 2017.

39. Waltham N, Schaffer J. Continuing aquatic assessment of wetlands with and without feral pig and cattle fence exclusion, Archer River catchment. Townsville Australia: TropWATER James Cook University, 2017 17/04.

40. Gongora J, Fleming P, Spencer PB, Mason R, Garkavenko O, Meyer J-N, et al. Phylogenetic relationships of Australian and New Zealand feral pigs assessed by mitochondrial control region sequence and nuclear GPIP genotype. Molecular Phylogenetics and Evolution. 2004;33(2):339–48.

41. Lopez J, Hurwood D, Dryden B, Fuller S. Feral pig populations are structured at fine spatial scales in tropical Queensland, Australia. PloS one. 2014;9(3):e91657.

42. Quinn GP, Keough MJ. Experimental design and data analysis for biologists: Cambridge University Press; 2002.

43. Clarke KR. Non-parametric multivariate analyses of changes in community structure. Australian Journal of Ecology. 1993;18:117–43.

44. De’Ath G. Multivariate regression trees: a new technique for modeling species–environment relationships. Ecology. 2002;83(4):1105–17.

45. Sheaves M, Johnston R. Ecological drivers of spatial variability among fish fauna of 21 tropical Australian estuaries. Marine Ecology Progress Series. 2009;385:245–60.

46. Bariyanga JD, Wronski T, Plath M, Apio A. Effectiveness of electro-fencing for restricting the ranging behaviour of wildlife: a case study in the degazetted parts of Akagera National Park. African zoology. 2016;51(4):183–91.

47. Arthington AH, Godfrey PC, Pearson RG, Karim F, Wallace J. Biodiversity values of remnant freshwater floodplain lagoons in agricultural catchments: evidence for fish of the Wet Tropics bioregion, northern Australia. Aquatic Conservation: Marine and Freshwater Ecosystems. 2015;25(3):336–52.

48. Barko VA, Herzog DP, O’Connell MT. Response of fishes to floodplain connectivity during and following a 500-year flood event in the unimpounded upper Mississippi River. Wetlands. 2006;26(1):244–57.

49. Chacin DH, Giery ST, Yeager LA, Layman CA, Langerhans RB. Does hydrological fragmentation affect coastal bird communities? A study from Abaco Island, The Bahamas. Wetlands ecology and management. 2015;23(3):551–7.

50. Robertson EP, Fletcher RJ, Austin JD. The Causes of Dispersal and the Cost of Carryover Effects for an Endangered Bird in a Dynamic Wetland Landscape. Journal of Animal Ecology. 2017.

51. Żmihorski M, Pärt T, Gustafson T, Berg Å. Effects of water level and grassland management on alpha and beta diversity of birds in restored wetlands. Journal of applied ecology. 2016;53(2):587–95.

52. Buelow CA, Baker R, Reside AE, Sheaves M. Nutrient subsidy indicators predict the presence of an avian mobile-link species. Ecological Indicators. 2018;89:507–15.

53. Ma Z, Cai Y, Li B, Chen J. Managing wetland habitats for waterbirds: an international perspective. Wetlands. 2010;30(1):15–27.

54. Swanson HK, Lysy M, Power M, Stasko AD, Johnson JD, Reist JD. A new probabilistic method for quantifying n-dimensional ecological niches and niche overlap. Ecology. 2015;96(2):318–24.

55. Jackson AL, Inger R, Parnell AC, Bearhop S. Comparing isotopic niche widths among and within communities: SIBER–Stable Isotope Bayesian Ellipses in R. Journal of Animal Ecology. 2011;80(3):595–602.

56. Pusey BJ, Kennard MJ, Douglas M, Allsop Q. Fish assemblage dynamics in an intermittent river of the northern Australian wet–dry tropics. Ecology of Freshwater Fish. 2018;27(1):78–88.

57. King A, Humphries P, Lake P. Fish recruitment on floodplains: the roles of patterns of flooding and life history characteristics. Canadian Journal of Fisheries and Aquatic Sciences. 2003;60(7):773–86.

58. Godfrey PC, Arthington AH, Pearson RG, Karim F, Wallace J. Fish larvae and recruitment patterns in floodplain lagoons of the Australian Wet Tropics. Marine and Freshwater Research. 2016:-. doi: https://doi.org/10.1071/MF15421.

59. Pusey BJ, Bird JR, Close AH, Arthington AH. Reproduction in three species of rainbowfish (Melanotaeniidae) in rainforest streams of north-eastern Queensland. Ecology of Freshwater Fish. 2001;10:75–87.

60. Stewart-Koster B, Olden J, Kennard M, Pusey B, Boone E, Douglas M, et al. Fish response to the temporal hierarchy of the natural flow regime in the Daly River, northern Australia. Journal of Fish Biology. 2011;79(6):1525–44.

61. Diaz RJ, Breitburg DL. The hypoxic environment. Fish physiology. 2009;27:1–23.

62. Butler B, Burrows DW. Dissolved oxygen guidelines for freshwater habitats of northern Australia. James Cook University, Townsville: Australian Centre for Tropical Freshwater Research (07/31), 2007 Contract No.: 07/31.

63. Collins GM, Clark TD, Rummer JL, Carton AG. Hypoxia tolerance is conserved across genetically distinct sub-populations of an iconic, tropical Australian teleost (Lates calcarifer). Conservation physiology. 2013;1(1):cot029.

64. Flint N, Pearson RG, Crossland MR. Reproduction and embryo viability of a range-limited tropical freshwater fish exposed to fluctuating hypoxia. Marine and Freshwater Research. 2018;69(2):267–76.

65. Gilmore KL, Doubleday ZA, Gillanders BM. Testing hypoxia: physiological effects of long-term exposure in two freshwater fishes. Oecologia. 2018;186(1):37–47.

66. James CS, Reside AE, VanDerWal J, Pearson RG, Burrows D, Capon SJ, et al. Sink or swim? Potential for high faunal turnover in Australian rivers under climate change. Journal of Biogeography. 2017.

67. Pusey BJ, Burrows DW, Kennard MJ, Perna CN, Unmack PJ, Allsop Q, et al. Freshwater fishes of northern Australia. Zootaxa. 2017;4253(1):1–104.

